# Jointly aligning cells and genomic features of single-cell multi-omics data with co-optimal transport

**DOI:** 10.1101/2022.11.09.515883

**Authors:** Pinar Demetci, Quang Huy Tran, Ievgen Redko, Ritambhara Singh

**Affiliations:** Center for Computational Molecular Biology, Brown University, RI, USA; Department of Computer Science, Brown University, RI, USA; Université Bretagne-Sud, CNRS, IRISA, Vannes, France; CMAP, Ecole Polytechnique, Palaiseau, France; Université Jean Monnet, CNRS, UMR 5516, Saint Etienne, France

**Keywords:** optimal transport, single-cell sequencing, multi-omics, regulatory genomics, data integration

## Abstract

The availability of various single-cell sequencing technologies allows one to jointly study multiple genomic features and understand how they interact to regulate cells. Although there are experimental challenges to simultaneously profile multiple features on the same single cell, recent computational methods can align the cells from unpaired multi-omic datasets. However, studying regulation also requires us to map the genomic features across different measurements. Unfortunately, most single-cell multi-omic alignment tools cannot perform these alignments or need prior knowledge. We introduce scootr, a co-optimal transport-based method, which jointly aligns both cells and genomic features of unpaired single-cell multi-omic datasets. We apply scootr to various single-cell multi-omic datasets with different types of measurements. Our results show that scootr provides quality alignments for unsupervised cell-level and feature-level integration of datasets with sparse feature correspondences (e.g., one-to-one mappings). For datasets with dense feature correspondences (e.g., many-to-many mappings), our joint framework allows us to provide supervision on one level (e.g., cell types), thus improving alignment performance on the other (e.g., genomic features) or vice-versa. The unique joint alignment framework makes scootr a helpful hypothesis-generation tool for the integrative study of unpaired single-cell multi-omic datasets.

**Available at**: https://github.com/rsinghlab/SCOOTR.

## 1 Introduction

Recent experimental developments have enabled us to measure various aspects of the genome, such as gene expression, chromatin accessibility, and methylation, at the single-cell resolution [1–4]. Jointly studying multiple genomic views can allow biologists to learn the rules of cell regulation by combining information about different genomic events. Although single-cell co-assay experiments can profile multiple measurement types on the same cell, they are only available for a few combinations of genomic signals [4], and can yield noisier data than single-omic experiments [5]. As a result, various computational methods [6–12] have been developed to successfully align single-cell datasets from non-co-assay (i.e. unpaired) experiments.

Among the existing unsupervised single-cell multi-omic alignment methods, optimal transport-based approaches [9–11] have shown state-of-the-art performance for aligning separately profiled (i.e. unpaired, non-co-assay) multi-omic datasets. Optimal transport solves for a probabilistic correspondence matrix between the data points of two input domains in order to match them in the most cost effective way possible [13]. A popular view of the problem is to imagine moving a pile of sand to fill in a hole through the least amount of work. One challenge in matching cells from multi-omic datasets is that this requires matching data points from different metric spaces. Existing methods [9–11] use Gromov-Wasserstein (GW) optimal transport [14, 15] for single-cell multi-omic alignment. GW optimal transport allows for relating data points from different metric spaces by comparing their intra-domain pairwise distances. Unfortunately, by computing intra-domain distances, it solely focuses on cell-cell mappings and shrouds the feature-feature mapping information of the input data. While obtaining cell-cell alignment is important, we also need to study the relationships between the features of different genomic measurements to understand how the input space of one measurement is optimally transformed into another. This study of regulatory relationships, therefore, requires the alignment of features. Such an alignment can also provide interpretations that guide researchers in learning about regulation at both cell and genomic feature level. Since, performing feature alignments is difficult using the GW optimal transport formulation, new computational approaches are needed to infer these alignments due to the high number of features and the complexity of their relationships.

Out of all existing single-cell alignment methods, only one method, bindSC [12], can (internally) perform feature-feature alignments. However, it requires some prior knowledge of their relationships using a gene activity matrix. This matrix is computed between gene expression features and the chromatin accessibility or methylation signals in the neighborhood of these genes. Therefore, the usability of bindSC is limited to measurement modalities with known relationships (usually with gene expression). Additionally, the default way of computing input gene activity matrices ignores most intergenic features in the chromatin accessibility and methylation modalities.

We introduce scootr (Figure 1), a co-optimal transport-based method [16] that simultaneously aligns both the cells and the features of unpaired single-cell multi-omic datasets in a measurement-agnostic manner. We apply scootr to diverse simulated and real-world single-cell datasets like – (1) a CITE-seq dataset with gene expression and epitope abundance measurements from single-cells, (2) a SNARE-seq dataset with chromatin accessibility and gene expression profiles, and (3) a multi-species single-cell RNA-seq dataset with gene expression measurements from bearded lizard pallium and mouse prefrontal cortex. The multi-species dataset is constructed from independent sequencing experiments, while the first two are co-assays with paired measurements. We use the ground-truth information on cell matches from these paired measurements to benchmark our cell-level alignments, and show that scootr is competitive with the current unsupervised cell-cell alignment methods. For feature-level alignments, we demonstrate with both simulated and real-world datasets that – (1) when feature correspondences are sparse (e.g., one-to-one mappings), such as in CITE-seq dataset, scootr yields high quality alignments without supervision, with 72% of the epitopes matching most strongly with the genes encoding them, and (2) when feature correspondences are dense (e.g., many-to-many mappings), such as in SNARE-seq, supervision on one level (e.g. cell-type alignments) improves the alignment on the other (e.g. features), or vice-versa, and yields up to ~ 80 – 100% accuracy on feature and cell-type matching tasks, respectively. This supervision at both cell and feature-level is uniquely possible due to our joint formulation.

**Figure 1:**
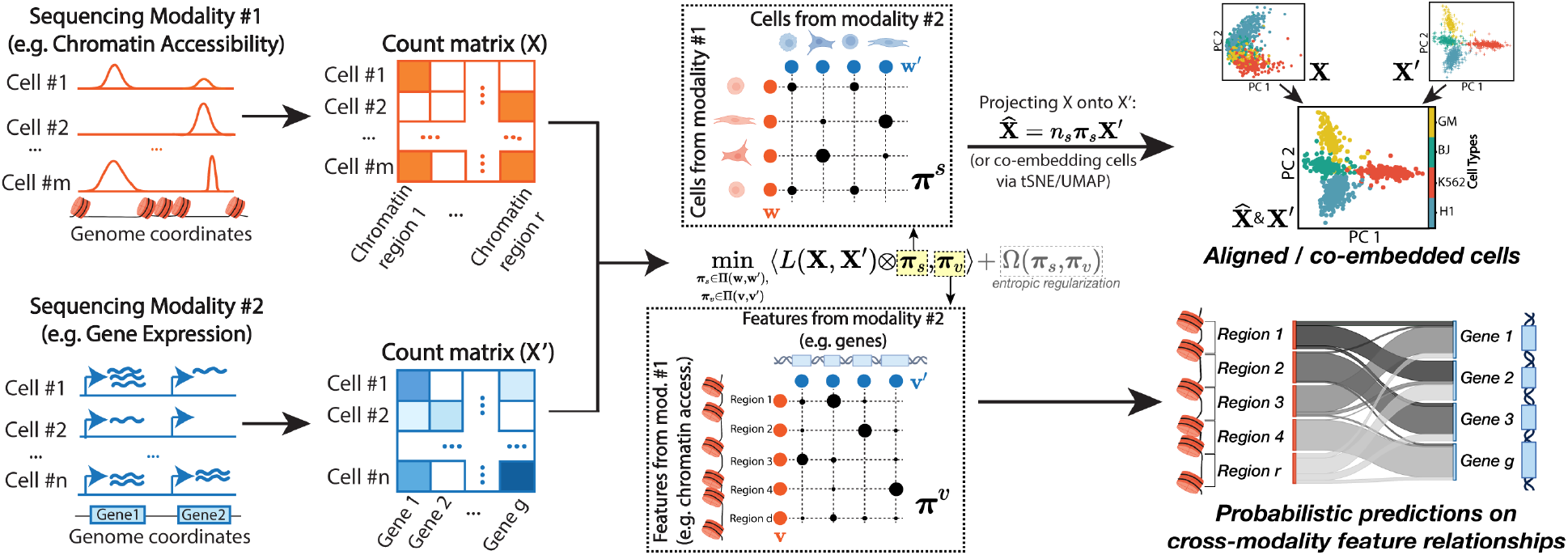
Overview of scootr on the SNARE-seq dataset. scootr takes in two count matrices, each from a different single-cell sequencing experiment. Given these unpaired single-cell datasets, scootr simultaneously solves for two probabilistic correspondence matrices: one between features, and one between cells across the two datasets. It returns the feature-feature coupling matrix for the user to investigate the correspondence probabilities. It uses the cell-cell coupling matrix to align the samples in the same space via barycentric projection or co-embedding via tSNE. Here, we visualize the alignment of the cells from SNARE-seq dataset via barycentric projection of the chromatin accessibility domain onto the gene expression domain, with data points colored by cell-type labels.

## 2 Method

Our method relies on optimal transport framework to align single-cell multi-omic datasets. We give a brief overview of optimal transport in Supplementary Section 5. Here, we first review the existing optimal transport-based methods to highlight our differences, then we introduce our framework.

### Notations

In what follows, we denote by 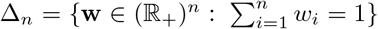 the simplex histogram with *n* bins. We use ⊗ for tensor-matrix multiplication, *i.e*., for a tensor **L** = (*L_i,j,k,l_*), the tensor-matrix multiplication **L** ⊗ **B** is the matrix (Σ_*k,l*_ *L_i,j,k,l_ B_k,l_*)_*i,j*_. We use 〈·,·〉 for the matrix scalar product associated with the Frobenius norm ‖ · ‖_*F*_. Finally, we write **1**_*d*_ ∈ ℝ^*d*^ for a *d*-dimensional vector of ones and denote all matrices by upper-case bold letters (*i.e*., **X**) or lower-case Greek letters (*i.e*., ***π***); all vectors are written in lower-case bold (*i.e*., **x**). We use the terms “coupling matrix” and “correspondence matrix” interchangeably.

### Related Previous Work

Previously, three optimal transport-based methods have been developed to integrate single-cell multi-omic datasets: SCOT [9], Pamona [10], and SCOTv2 [11]. All three methods rely on Gromov-Wasserstein optimal transport to align cells from single-cell datasets that measure different genomic features. Optimal transport formulation solves for a matrix of correspondence probabilities that will transform and align the datasets in a way that leads to minimal cost. However, defining a cost metric over samples from different metric spaces (i.e. from different measurement modalities) is challenging. Gromov-Wasserstein (GW) distance allows for the comparison of distributions in different metric spaces by comparing pairwise distances between the samples across these domains. This also preserves the local neighborhood geometry when moving data points between domains.

Given two datasets (or single-cell measurements) represented by matrices **X** = [**x**_1_,…, **x**_*n*_]^*T*^ ∈ ℝ^*n*×*d*^ and 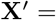 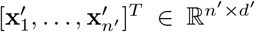, we let 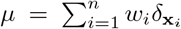 and 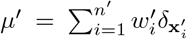 be two empirical distributions related to samples, where **x**_*i*_ ∈ ℝ^*d*^ and 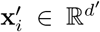. Here, 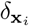 is the Dirac measure. We refer in the following to **w** = [*w*_1_,…, *w_n_*]^⊤^ ∈ Δ_*n*_ and 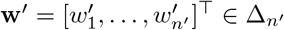 as sample weights vectors that both lie in the simplex. For the Gromov-Wasserstein formulation, one first computes pairwise distance matrices **D^x^** and **D^x′^** with 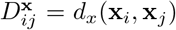 and 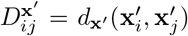 for some distances *d*_**x**_ and *d*_**x′**_ [14].Then, for a given cost function *L* : ℝ × ℝ → ℝ, defined over the distance matrices, the discrete GW distance solves the following optimization problem:

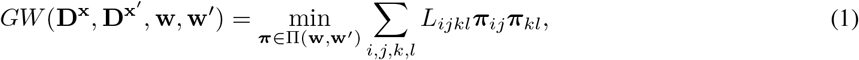

where 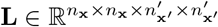 is the fourth-order tensor defined by 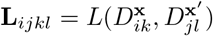 and Π(·, ·) is the set of linear transport constraints defined as:

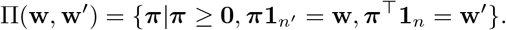

Intuitively, 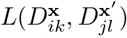 captures how transporting **x**_*i*_ onto 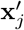 and **x**_*k*_ onto 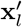 would distort the original distances between **x**_*i*_ and **x**_*k*_ and between 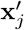 and 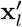. The major difference between SCOT, SCOTv2, and Pamona are the regularization terms they use in the Gromov-Wasserstein formulation. All three alignment methods use *L*(*x, y*) = (*x* − *y*)^2^ and use the cell-cell coupling matrix to align cells in a joint space.

### Co-Optimal transport

While the intra-domain distances used in the GW-based methods often provides richer representations of datasets, it also comes with several important drawbacks. First, GW requires finding the right similarity measure for a given dataset, which may be hard and computationally costly in practice when no expert knowledge is available. Second, GW exhibits invariances which, as shown before [16], may lead to less precise sample matchings in some applications. Most importantly, GW-based optimal transport drops the features of the datasets after computing intra-domain distances, with no opportunity for their alignment.

In order to address these drawbacks, Redko *et al*. [16] proposed a new OT problem termed Co-Optimal transport (COOT) that aligns samples and features of the two datasets in the original space. We define COOT formulation using the same notation as before. In addition, we now also define weights for the features that are stored in vectors **v** ∈ Δ_*d*_ and **v′** ∈ Δ_*d*′_.

Co-Optimal Transport then reads as follows:

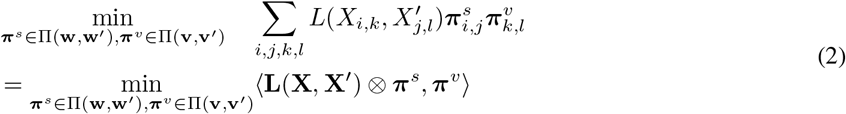

where **L** is the tensor of all pairwise divergences between the elements of the matrices **X** and **X′**. Intuitively, in case of images, for instance, this tensor will consists of all differences between individual pixels of the two images, while for GW those will be the differences between the similarities of all pixels within the image.

As explained above, (2) optimizes over a coupling ***π***^*s*^ between samples and a coupling ***π***^*v*^ between features of the two datasets. The intuition behind optimizing over two couplings is as follows: while the samples coupling ***π***^*s*^ has the same meaning and acts in the same way as the coupling returned by GW, the feature coupling ***π***^*v*^ returned by COOT aims to align the distributions over the features that describe the two datasets. In the simplest case, this may correspond to finding a permutation of the features that leads to the sample alignment of the smallest cost. In more complex settings, such as, for instance, when aligning images of different size, this may correspond to finding a spatial transformation that up/downscales the image following the least effort principle. We write COOT(**X**, **X′**) to denote the objective value of the optimization problem (2) when uniform weight vectors are used (as done for GW before).

COOT can be extended to include entropic regularization [17] term, that is popular in OT community, as follows:

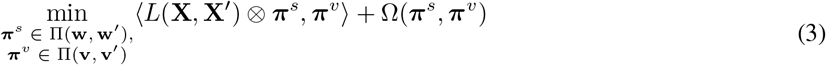

where for *ϵ*_1_, *ϵ*_2_ > 0, the regularization term writes as Ω(***π***^*s*^, ***π***^*v*^) = *ϵ*_1_*H*(***π***^*s*^ |**ww**^′*T*^)+*ϵ*_2_*H*(***π***^*v*^ |**vv**^*′T*^) with 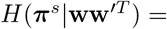 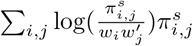 being the relative entropy. Note that similarly to OT [17] and GW [15], adding the regularization term can lead to a more robust and fast estimation of couplings but prevents them from being sparse.

### Single-cell CO-Optimal TRansport (SCOOTR)

Our proposed method, scootr (Figure 1), directly takes the normalized count matrices of the single-cell measurements (**X** and **X′**) as inputs. Unlike previous OT-based methods, we do not require to calculate the pairwise similarities, thus reducing the dependence on the choice of similarity function and its associated hyperparameters. We set the marginal distributions over samples, **w**, **w′** and features **v**, **v′** as uniform distributions assuming no prior knowledge about the datasets, however, we allow the users to customize the weights in these distributions if they have any prior information (e.g. scaling up the weight of features with regulatory significance that are expected to match with many other features). Given the input datasets and the marginal distributions, scootr solves for cell-cell and feature-feature correspondence matrices using the co-optimal transport formulation:

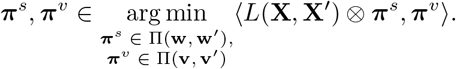

where 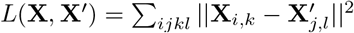 for cells *i, j* and features *k, l*.

The optimization is carried out with a block coordinate descent procedure proposed by Redko *et al* [16], which is detailed in the Supplementary Algorithm 1. This procedure alternates between optimizing the two matrices in each optimization iteration.

After computing the coupling matrices, scootr aligns the cells of the two datasets in a common space using the cell-cell alignment probabilities given in (***π***^*s*^) either by barycentric projection 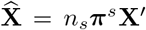, or by co-embedding them using t-SNE (described in Supplementary Algorithm 1). scootr returns the feature-feature coupling matrix for the user to further investigate the feature relationships. However, we provide an interface with ReMap Atlas of Regulatory Regions API. The probabilities in the coupling matrix can allow users to rank correspondences and prioritize downstream analyses.

A useful property of the co-optimal transport formulation is that one can provide a weak supervision when solving for ***π***^*s*^ and ***π***^*v*^ by scaling the costs of matching samples/features that should not be matched together. We do this by multiplying the cost matrix ***L*** with a supervision matrix ***D*** that user provides in inputs. This matrix can be provided for either the sample or the feature alignments. If it’s provided for the feature-level alignments, for example, it is only used in the optimization step for the feature-feature coupling matrix in the block coordinate descent (Supplementary Algorithm 1). The entries of the supervision matrix are expected to range between 0 and 1. For example, an entry for 0 for the row *i* and column *j* in the feature-level supervision matrix removes the cost associated with aligning the feature *i* and feature *j* of the two input datasets, respectively. This is particularly useful when the underlying feature relationships are expected to be less sparse and more complex than 1-1 matches. We will show that this unique feature of scootr is valuable when dealing with single-cell measurements.

## 3 Experimental Setup

We first show that scootr can yield cell-cell alignment results on par with the existing single-cell multi-omic alignment methods. Then, we demonstrate its ability to also simultaneously align the features from different genomic modalities, using simulated and real-world single-cell sequencing datasets.

### 3.1 Datasets

When choosing datasets, we follow our main baselines – the existing optimal transport-based single-cell alignment methods [9–11] and bindSC [12] – and curate a similar set of simulated and real-world datasets for a comparable benchmarking. We use the datasets with ground-truth information on 1-1 cell pairings for evaluating cell-level alignment performance. Similarly, we use the datasets with some prior information on feature correspondences for evaluating feature-level alignment performance.

#### Datasets for cell-cell alignment benchmarking

We use four simulated datasets and three real-world single-cell multi-omic datasets to benchmark our cell alignment performance. Three of the simulated datasets have been generated by Liu *et al*. [6] by non-linearly projecting 600 samples from a common 2-dimensional space onto different 1000- and 2000-dimensional spaces with 300 samples in each. In the first simulation, the data points in each domain form a bifurcating tree structure that is commonly seen in cell populations undergoing differentiation. The second simulation forms a three dimensional Swiss roll. Lastly, the third simulation forms a circular frustum that resembles what is commonly observed when investigating cell cycle. These datasets have been previously used for benchmarking by other cell-cell alignment methods [6–10]. We refer to these datasets as “Sim 1”, “Sim 2”, and “Sim 3”, respectively. We include a fourth simulated dataset that has been generated by [9] using a single-cell RNA-seq data simulation package in R, called Splatter [18]. We refer to this dataset as “Synthetic RNA”. This dataset includes a simulated gene expression domain with 50 genes and 5000 cells divided across three cell-types, and another domain created by non-linearly projecting these cells onto a 500-dimensional space. As a result of their generation schemes, all simulated datasets have ground-truth 1-1 cell correspondence information.

Additionally, we include three real-world sequencing datasets. To have ground-truth information on cell correspondences for evaluation, we choose three co-assay datasets which have paired measurements on the same individual cells: an scGEM dataset [19], a SNARE-seq dataset [1], and a CITE-seq dataset [2]. These first two datasets have been used by existing single-cell alignment methods, including the ones employing optimal transport [6–11], while the last one was included in the evaluations of bindSC [12]. The scGEM dataset contains measurements on gene expression and DNA methylation states of 177 individual cells from human somatic cell population undergoing conversion to induced pluripotent stem cells (iPSCs) [19]. We accessed the pre-processed count matrices for this dataset through the MATCHER repository ^1^. The SNARE-seq dataset contains gene expression and chromatin accessibility profiles of 1047 individual cells from a mixed population of four cell lines: H1(human embryonic stem cells), BJ (a fibroblast cell line), K562 (a lymphoblast cell line), and GM12878 (lymphoblastoid cells derived from blood) [1]. We access their count matrices on Gene Expression Omnibus, with the accession code GSE126074. Finally, the CITE-seq dataset has gene expression profiles and epitope abundance measurements on 25 antibodies from 30,672 cells from human bone marrow tissue [2]. The count matrices for this dataset were downloaded from the Seurat website ^2^.

#### Datasets for feature-feature alignment benchmarking

We assess feature-level alignment performance on real-world single-cell multi-omic datasets with some ground-truth correspondence information between the features. Among the three real-world datasets described above, we have ground-truth information on the CITE-seq dataset, where we know which genes from the gene expression domain encode the 25 antibodies from the antibody abundance domain. We use these 1-1 correspondences to evaluate our feature alignments. Since we do not have such reliable ground-truth feature correspondence information for SNARE-seq and scGEM datasets, we use a novel computational tool called CellOracle [20]. This tool has been developed to infer regulatory networks jointly from single-cell chromatin accessibility and gene expression data. For the SNARE-seq dataset, we use CellOracle to construct gene regulatory networks. We take the chromosomal region of transcription factors and genomic elements that a gene is connected to in this network as its probable feature correspondences, and compare our alignments against these (more detail in Section 3.2). We are unaware of the existence of such tools for single-cell methylation data. As a result, we do not include scGEM dataset in our feature-level alignment performance benchmarking. Instead, we add a new real-world dataset with a need for single-cell alignment. This dataset contains unpaired single-cell gene expression profiles from mouse prefrontal cortex[21], and the bearded lizard pallium [22]. It has been curated with data from separately conducted experiments. Here, we have information on paralogous genes across the two species, as well as relevant cell types.

In addition to these three real-world sequencing datasets, we simulate a new set of multi-omic data with varying levels of sparsity in underlying feature-level correspondences. Our goal for including these simulations is to investigate the effect of correspondence sparsity on alignment performance. We follow the simulation set-up by Zhang *et al* [23], which modifies a single-cell RNA-seq simulation method, SymSim [24], to also simulate scATAC-seq count data based on a gene-chromosomal region relationship matrix ^3^. We simulate 500 cells with 50 genes in the gene expression modality, and 1000 chromosomal regions in the chromatin accessibility modality. We randomly generate five gene-to-chromosomal region correspondence matrices with uniform 1-2 (sparse), 1-4, 1-6, 1-8, and 1-10 (dense) matches. We generate five multi-omic datasets using these ground-truth correspondence matrices.

### 3.2 Evaluation Criteria

#### Cell-cell alignment evaluation

When evaluating cell-cell alignments, we use a metric previously used by other single-cell multi-omic integration tools [6–12] called “fraction of samples closer than the true match” (FOSCTTM). For this metric, we compute the Euclidean distances between a fixed sample point and all the data points in the other domain. Then, we use these distances to compute the fraction of samples that are closer to the fixed sample than its true match, and then average these values for all the samples in both domains. This metric measures alignment error, so the lower values correspond to higher quality alignments.

#### Feature-feature alignment evaluation

To assess feature-feature alignment performance, we investigate the accuracy of feature correspondences recovered. We mainly use three real-world datasets for this task - CITE-seq, SNARE-seq, and the cross-species scRNA-seq datasets. Due to the versatility of the genomic measurements in these datasets, we follow a different procedure for each to define “ground-truth” feature correspondences to compute the accuracy.

For the CITE-seq dataset, we expect the feature correspondences to recover the relationship between the 25 antibodies and the genes that encode them. To investigate this, we simultaneously align the cells and features of the two modalities using the 25 antibodies and 25 genes in an unsupervised manner. We compute the percentage of 25 antibodies whose strongest correspondence is their encoding gene.

For SNARE-seq dataset, we start by pre-processing the dataset to prune features. The original dataset contains 18,666 genes and 1,136,771 chromosomal regions in their respective modalities. We select the top 1000 most variable genes using the `FindVariableFeatures` function of Seurat with its default parameters. We also select the top 2500 chromosomal regions in the chromatin accessibility domain using the `FindTopFeatures` function of Signac [25]. With these, we construct a gene regulatory network with CellOracle [20] using both domains. Then, for each gene, we identify the chromosomal regions of its regulators from the regulatory network, using the human reference genome GRCh38/hg38. We expect the genes in the gene expression modality to be matched with at least one of the chromosomal regions overlapping with each regulator’s genomic coordinates, and compute the accuracy over all genes accordingly.

For the cross-species RNA-seq dataset, we expect alignments between the cell-type annotations common to the mouse and lizard datasets, namely: excitatory neurons, inhibitory neurons, microglia, OPC (Oligodendrocyte precursor cells), oligodendrocytes, and endothelial cells. For this dataset, we generate cell-label matches by averaging the rows and columns of the cell-cell alignment matrix yielded by scootr based on these cell annotation labels. We compute the percentage of these six cell-type groups that match as their strongest correspondence.

### 3.3 Baselines

For the cell-cell alignment evaluation, we consider the following unsupervised single-cell multi-omic integration methods, MMD-MA [6], UnionCom [7], SCOT [9, 11], Pamona [10], and bindSC [12]. Among these, SCOT and Pamona are optimal transport-based methods; they both use GW optimal transport with different relaxations. We note that unlike other baselines, bindSC could be considered a weakly supervised method since it requires a gene activity matrix as an input. For feature-feature alignment benchmarking, bindSC remains our only baseline since the other integration methods only perform alignment on the cell level. Although bindSC does not return the final feature-level correspondence matrix to the user, it does return the relationship between each feature and the computer intermediary factors. By multiplying these matrices, we are able to obtain a feature-level correspondence matrix. For all methods, we set a grid of hyperparameter combinations and choose the best performing combination for each dataset. For scootr, we consider the EMD (for *ϵ*_1,2_ = 0) and Sinkhorn algorithms for each OT subproblem, and entropic regularization strength for Sinkhorn taking values in 10^−5^,…, 10^4^. The hyperparameter combinations considered for the baselines are described in Supplementary Section 5.

## 4 Results

### 4.1 scootr gives comparable performance to existing cell-cell alignment methods

We integrate four simulated and three real-world co-assayed datasets to compare scootr’s cell-cell alignment performance with the existing unsupervised alignment methods. Table 1 shows the cell alignment errors yielded by scootr and each of our baselines, as measured by the average FOSCTTM metric. Among the baselines considered on this table, we draw special attention to SCOT and Pamona, which are GW optimal transport-based methods. These methods align cells by comparing pairwise distances between cells, discarding their features. By benchmarking against them, we show that aligning cells based on raw features can lead to competitive results. We observe that while SCOT tends to outperform scootr on co-assay datasets, scootr yields alignments of similar quality for scGEM and SNARE-seq datasets and outperforms SCOT on simulation datasets. scootr also outperforms Pamona on aligning cells from the SNARE-seq dataset while yielding a very similar alignments for the scGEM and CITE-seq dataset. Cell alignments yielded by scootr for the CITE-seq and SNARE-seq datasets are visualized in Figure 2B and Figure 3B (original domains before alignment visualized in A) and the alignment for scGEM dataset is visualized in Figure S1. We obtain these alignments by projecting one domain (gene expression for CITE-seq, chromatin accessibility for SNARE-seq, and DNA methylation for scGEM) onto the other domain using barycentric projection. Overall, they show that the cell-type clusters are preserved upon alignment by scootr.

**Table 1:** Benchmarking scootr against existing single-cell multi-omic alignment methods on sample-level (cell) alignment. The numbers indicate the average fraction of samples a sample is aligned closer to than its true match (FOSCTTM). This metric measures alignment error and lower values indicate higher quality alignments. Here, we highlight the best results in bold and underline the second best ones.

|  | S1 | S2 | S3 | Synthetic RNA | sc-GEM | SNAREseq | CITE-seq |
| --- | --- | --- | --- | --- | --- | --- | --- |
| **SCOT** | 0.0866 | 0.0216 | <u>0.0084</u> | 0.000071 | **0.198** | **0.15** | **0.098** |
| **UnionCom** | 0.083 | 0.0157 | 0.152 | 0.038 | 0.2096 | 0.265 | 0.169 |
| **MMD-MA** | 0.1244 | 0.0327 | 0.01246 | 0.112 | <u>0.2014</u> | **0.15** | <u>0.113</u> |
| **Pamona** | <u>0.078</u> | <u>0.011</u> | **0.0082** | <u>0.00041</u> | 0.204 | 0.2217 | 0.116 |
| **bindSC** | N/A* | N/A* | N/A* | N/A* | 0.2151 | 0.273 | 0.130 |
| **SCOOTR** | **0.07** | **0.004** | 0.0088 | **0** | 0.206 | <u>0.153</u> | 0.132 |

**Figure 2:**
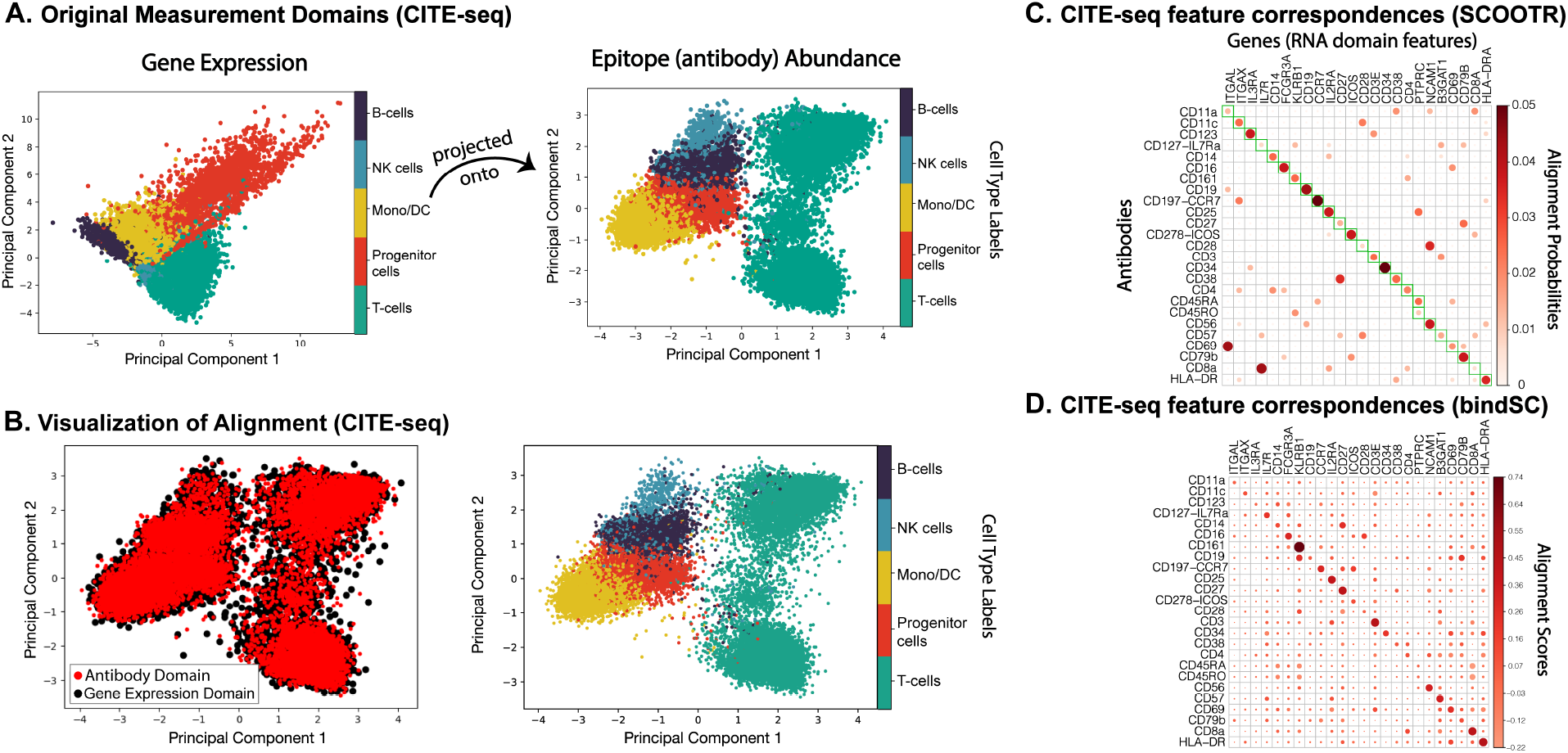
Cell-cell and feature-feature alignment results on CITE-seq dataset. **A** visualizes the original domains – antibody abundance, and gene expression, respectively –, following dimensionality reduction with 2D principal component analysis (PCA). **B** visualizes the aligned domains, after the gene expression domain has been projected onto the antibody abundance domain via barycentric projection. **C** Feature alignment probabilities recovered by scootr. The green boxes along the diagonal indicate the “ground-truth” correspondences we expect to see between the antibodies and their encoding genes. **D**. The feature alignment probabilities recovered by bindSC.

**Figure 3:**
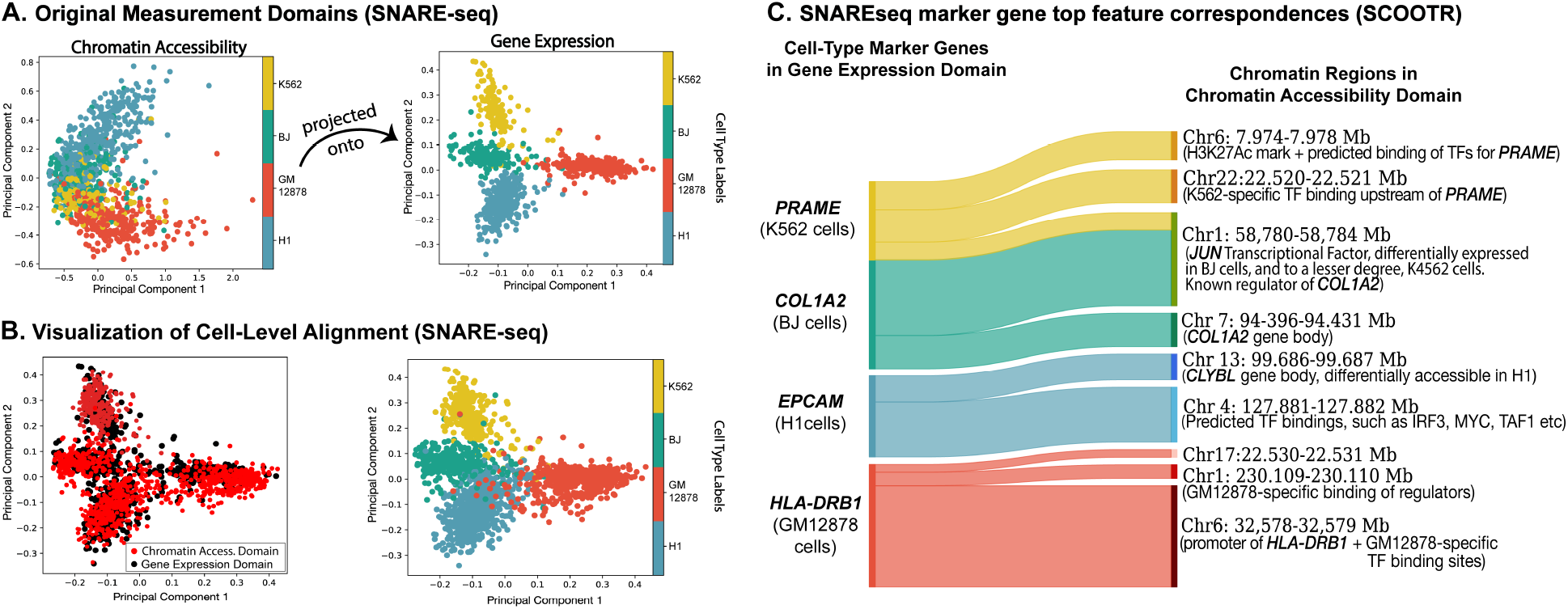
Cell and feature alignment results on SNARE-seq dataset. **A** visualizes the original domains – chromatin accessibility, and gene expression, respectively, following dimensionality reduction with 2D principal component analysis (PCA). **B** visualizes the aligned domains, after the chromatin accessibility domain has been projected onto the gene expression domain domain via barycentric projection. **C** Sankey plot visualizing the top chromatin accessibility feature correspondences recovered for the cell-type marker genes. These correspondences include the chromosomal regions of the marker genes and regions with predicted cell-type specific transcriptional factor (TF) binding.

Among our baselines, SCOT has a heuristic for self-tuning hyperparameters by tracking the Gromov-Wasserstein distance between aligned datasets. We demonstrate in Figure S2 that scootr can also perform hyperparameter tuning in a similar way. We observe that lowe values of co-optimal transport distances tend to correspond to lower average FOSCTTM values (alignment error). Therefore, despite replacing the Gromov-Wasserstein formulation, we are able to retain the approximate self-tuning procedure of the other optimal transport-based methods.

### 4.2 scootr can simultaneously align features across single-cell multi-omic datasets

Next, we investigate the feature alignments yielded by scootr. We demonstrate that while scootr can recover both cell-level and feature-level correspondences in an unsupervised manner for datasets with highly sparse correspondences (CITE-seq), for others with dense feature relationships (e.g. SNARE-seq), it greatly benefits from receiving some supervision. In what follows, we first present our results on unsupervised alignment of the CITE-seq dataset, and then demonstrate that providing supervision on cell-level alignments can improve feature-level alignment performance and vice versa. For the feature alignment task, bindSC remains our only baseline because the other single-cell alignment methods only perform cell-cell alignment.

#### 4.2.1 Unsupervised alignment

Among the real-world datasets we use, CITE-seq has underlying 1-1 correspondences between the antibodies and their encoding genes. We expect scootr to recover them when we align 25 antibodies with the corresponding 25 genes (while simultaneously aligning cells) in its unsupervised setting. We present the feature correspondence matrix scootr yields in 2C. The rows and columns of this matrix are ordered such that the expected ground-truth correspondences are along the diagonal (marked by green squares). Note that the row and column probabilities add up to the weights from marginal distributions initialized in the beginning of the optimization. We observe that all antibodies are matched to their corresponding genes with a non-zero probability of correspondence and 18 of them (~70%) have the strongest correspondence probability with their encoding gene. Compared to the feature correspondence matrix we receive from bindSC (Figure 2D), we yield a sparser correspondence matrix while still correctly aligning a similar number of antibodies with their encoding genes. Figure 2B shows the cell-level alignments we receive from this run, which demonstrates that scootr simultaneously recovers quality cell and feature alignments for CITE-seq dataset.

#### 4.2.2 Cell-level supervision for improved feature alignment

Despite the success of unsupervised alignment on CITE-seq dataset, we observe that when the underlying correspondences are dense with many-to-many matches, recovering them in an unsupervised fashion proves to be more challenging. We demonstrate this on simulated multi-omic datasets (scRNA-seq and scATAC-seq), which have been generated with varying levels of feature correspondence sparsity (1-2, 1-4,.. 1-10) between 50 simulated genes (in the scRNA-seq domain) and 100 chromosomal regions (in the scATAC-seq domain). Figure 4B demonstrates that scootr’s feature correspondence recovery performance decreases with lower levels of correspondence sparsity. Similarly, the dense feature correspondences in SNARE-seq dataset make it more difficult to recover them without any guidance. However, in real-world applications, we can reasonably expect some level of prior information to be available on the samples, namely, cell-type annotations, for at least a subset of the cells. In most cases, upon obtaining sequencing measurements, biologists use marker genes to find different cell-type groups. Although it is expected to observe some mismatch in cell type annotations between different genomic views, we demonstrate in Table 2 that the performance of scootr improves even with weak supervision. Our unique joint alignment formulation provides the ability to perform this weak supervision at both sample and feature level. In this table, we use varying proportions (0, 20, …100%) of the cell-type annotations to provide supervision on the cell-level to assist with the feature-level alignments. As described in Section 2, we create a supervision matrix, which removes the cost of aligning two cells if they belong to the same cell type. In the 100% supervised setting, we use all of the cell type annotations to create this supervision matrix; whereas in the 20% supervised setting, we only use 20% of the cell type annotations. To obtain a ground-truth on correspondences, we use CellOracle [20] to infer a regulatory network using both the gene expression and the chromatin accessibility data. We expect to recover the correspondences between genes and at least one segment of the chromosomal regions corresponding to each of their regulators (described in Section 3.2). We visualize examples of chromatin accessibility to gene expression correspondences yielded by scootr for the cell-type marker genes in Figure 3(C).

**Table 2:**
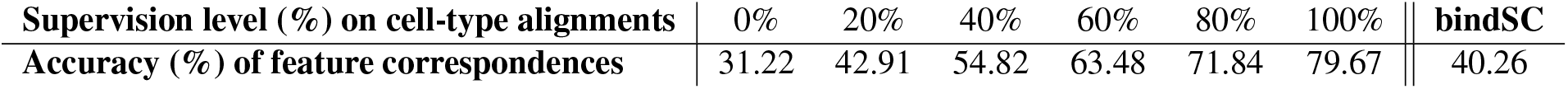
Feature alignment performance on SNARE-seq with increasing supervision on cell-type alignments.

| Supervision level (%) on cell-type alignments | 0% | 20% | 40% | 60% | 80% | 100% | bindSC |
| --- | --- | --- | --- | --- | --- | --- | --- |
| Accuracy (%) of feature correspondences | 31.22 | 42.91 | 54.82 | 63.48 | 71.84 | 79.67 | 40.26 |

**Figure 4:**
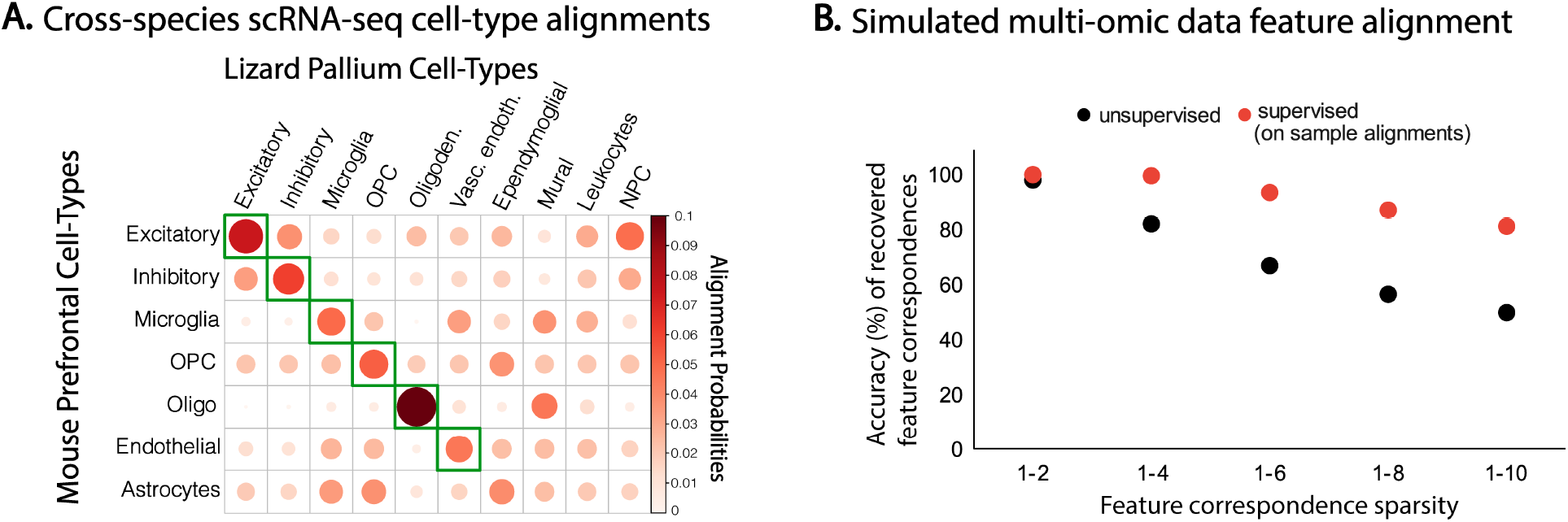
Cell and feature alignment results on SNARE-seq dataset. **A** visualizes the cell-type alignment probabilities across the mouse prefrontal cortex and bearded lizard pallium. The larger and darker the points are, the higher the alignment probability. Green boxes indicate the common cell-type annotations. **B** plots the feature-feature alignment accuracy across varying levels of sparsity in ground-truth correspondences in scootr’s unsupervised setting (black) and supervised setting with full supervision provided on the sample-sample alignments (red).

Furthermore, we look into biological annotations of the matching chromatin accessibility regions on UCSC Genome Browser [26] with annotations from JASPAR Transcription Factor Binding Site Database [27] and ReMap Atlas of Regulatory Regions [28]. We observe that the marker genes are matched with their chromosomal region or the regions associated with relevant transcription factor binding sites. For example, the strongest correspondence of *COL1A2*, the marker gene of BJ cell-line, is in the chromosomal region of *JUN*, which is a transcription factor identified to be differentially expressed in BJ cells, and to a lesser level, K562 cells [1]. Its second strongest match is a region within its own chromosomal region. Similarly, the marker gene for the GM12878 cell line, *HLA-DRB1*, is most strongly matched with a region upstream of its own genomic region, along with predicted GM12878-specific transcriptional binding sites. We detail these correspondences in Supplementary Materials Section 5. Overall, we see that scootr can recover biologically meaningful feature correspondences with supervision on the cell-type alignments. This can allow biologists to generate hypotheses on regulatory relationships between genomic features from different measurements, and prioritize further investigations based on the ranking of correspondence probabilities.

#### 4.2.3 Feature-level supervision for improved cell alignment

Here, we demonstrate that supervision on the feature-level alignments improves cell-level alignment quality. For this, we align the gene expression data obtained from the brain tissue of two different species, namely, mouse prefrontal cortex and bearded dragon lizard pallium. Since these are separately profiled datasets, we do not expect to find 1-1 correspondences between the cells; however, we have prior information on relatedness of cell-type labels (Section 3.2). Here, we provide feature-level supervision to guide scootr to recover these cell-type relationships. Coming from different species, these datasets do not have the exact same set of features, however, they share 10,816 paralogous genes. We create a supervision matrix, similar to the case in SNARE-seq, to guide the alignments between paralogous genes across the two datasets and investigate the alignments yielded for the cells. We average the cell-cell alignment probabilities across cell types to get the cell-type alignments. Figure 4A demonstrates the cell-type alignments we receive from fully supervised case, and the Table 3 presents the cell-type alignment accuracy with varying levels (0%, 20%, …, 100%) of feature alignment supervision. Here, accuracy is calculated as a percentage of correspondences recovered among all the expected correspondences (marked by the green boxes in Figure 4A). Similarly to cell alignment, we see that supervision on feature alignments increases cell-level alignment accuracy. So, when validation data is present on feature relationships, they can be used to obtain higher quality cell-cell alignments using scootr. Additionally, this application demonstrates that scootr can also be used to relate cell clusters from different datasets.

**Table 3:** Cell-type alignment performance on cross-species RNA-seq dataset with increasing supervision on paralogous gene alignments.

| Supervision level (%) on cell-type alignments | 0% | 20% | 40% | 60% | 80% | 100% | bindSC |
| --- | --- | --- | --- | --- | --- | --- | --- |
| Accuracy (%) of feature correspondences | 50.00 | 50.00 | 66.67 | 66.67 | 83.34 | 100.00 | 66.67 |

## 5 Discussion

The majority of the existing single-cell multi-omic alignment methods solely align cells. Our proposed method scootr jointly aligns both the cells and the features of single-cell multi-omic datasets, allowing researchers to study potential relationships between different views of the genome. We intend scootr to be a hypothesis-generation tool for biological scientists. The correspondence probabilities that scootr yields can be used to rank the predicted cross-measurement relationships, allowing scientists to prioritize downstream investigations accordingly when studying regulatory interactions.

Unlike scootr, Gromov-Wasserstein optimal transport-based alignment methods use dissimilarity kernels (based on pairwise sample distances) for alignment, which appears to show a slight advantage in cell-cell alignment performance, possibly because these kernels give a less noisy view of the data. However, these methods cannot align features since dissimilarity kernels discard them. On the other hand, scootr directly uses the count matrices, avoiding this downside. Therefore, the small performance trade-off in scootr’s cell alignments comes with the gain of jointly aligning features and also supporting the alignment of one level (e.g., cells) with supervision on the other (e.g., features). This formulation is beneficial for various multi-omics studies.

Single-cell alignment methods require validation data on cell-cell correspondence to tune the hyperparameters. However, such information is unlikely to be present in real-world cases when datasets are separately sequenced. Although both SCOT and scootr perform self-tuning by tracing optimal transport cost, the lowest cost does not always correspond to the best alignment, and the quality of self-tuning can vary between datasets. If prior information other than cell-cell alignment validations is present –such as the paralogous genes in the case of cross-species alignment experiments or the cell-type annotations in SNARE-seq experiments–, using these could lead to better alignments in some datasets compared to self-tuning. Our experiments demonstrate that even partial supervision leads to improvement in alignment performance.

For the feature-level alignments, neural-network-based formulations of co-optimal transport could potentially allow one to account for more complex relationships. However, maintaining feature-level interpretability in the coupling matrix becomes more challenging in this formulation. As future work, we are investigating an interpretability-preserving neural formulation. Additionally, it might be possible to set the marginal distributions over cells and features based on common biological knowledge. For example, when aligning gene expression data and chromatin accessibility data, one might scale the weights of chromosomal regions corresponding to common transcription binding sites based on existing databases, guiding the algorithm to align these with more genes. Similarly, one could scale the weights of the cells based on prior clustering without the need for cell-type annotations. Our future work will compare such approaches to the unsupervised and supervised cases presented here. In the meantime, we allow users to customize marginal distributions when running scootr. Overall we demonstrate that scootr is a competitive single-cell multi-omic data integration method that can help generate hypotheses for genomic feature relationships when jointly studying multiple single-cell datasets.

## Acknowledgements

We thank Rebecca Santorella, Rémi Flamary and Nicolas Courty for helpful discussions.

## Funding

Ritambhara Singh’s and Pinar Demetci’s contribution to the work is supported by NIH award R35HG011939.

## Supplementary Materials

### Background on Optimal Transport

Optimal transport is a mathematical framework for relating probability distributions or discrete measures to one another. Here, we focus on discrete measures because we work with sequencing datasets that contain empirical measurements on a finite set of samples. Consider two datasets in ℝ^*d*^ with *n* and *n*′ data points in each, represented by matrices **X** = [**x**_1_,…, **x**_*n*_]^⊤^ ∈ ℝ^*n*×*d*^ and 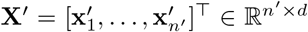. We let 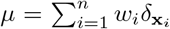 and 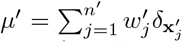 be two empirical distributions related to their samples. Here 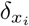 is the Dirac measure and we refer in the following to **w** = [*w*_1_,…, *w_n_*]^⊤^ ∈ Δ_*n*_ and 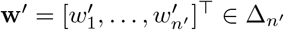 as sample weights vectors that both lie in the simplex.

Given a cost function 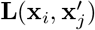 that describes how “expensive” it is to match one data point (**x**_*i*_) with another 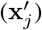 across the two datasets, Kantorovich formulation of optimal transport sets out to find an optimal coupling ***π*** that attains:

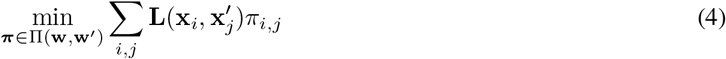

Here, the coupling ***π*** holds the alignment probabilities between each pair of data points across the two datasets to optimally transform one into the other. Most of the practical applications of optimal transport includes an entropic regularization over the coupling matrix to split the alignment probabilities across multiple samples and also to make the optimization computationally more efficient. For more detailed background on optimal transport, we refer readers to Villani, 2008 [29] (for theory), and Peyré and Cuturi (2019) [13] (for computational aspects).

### Methods: Algorithms

**Algorithm 1:**
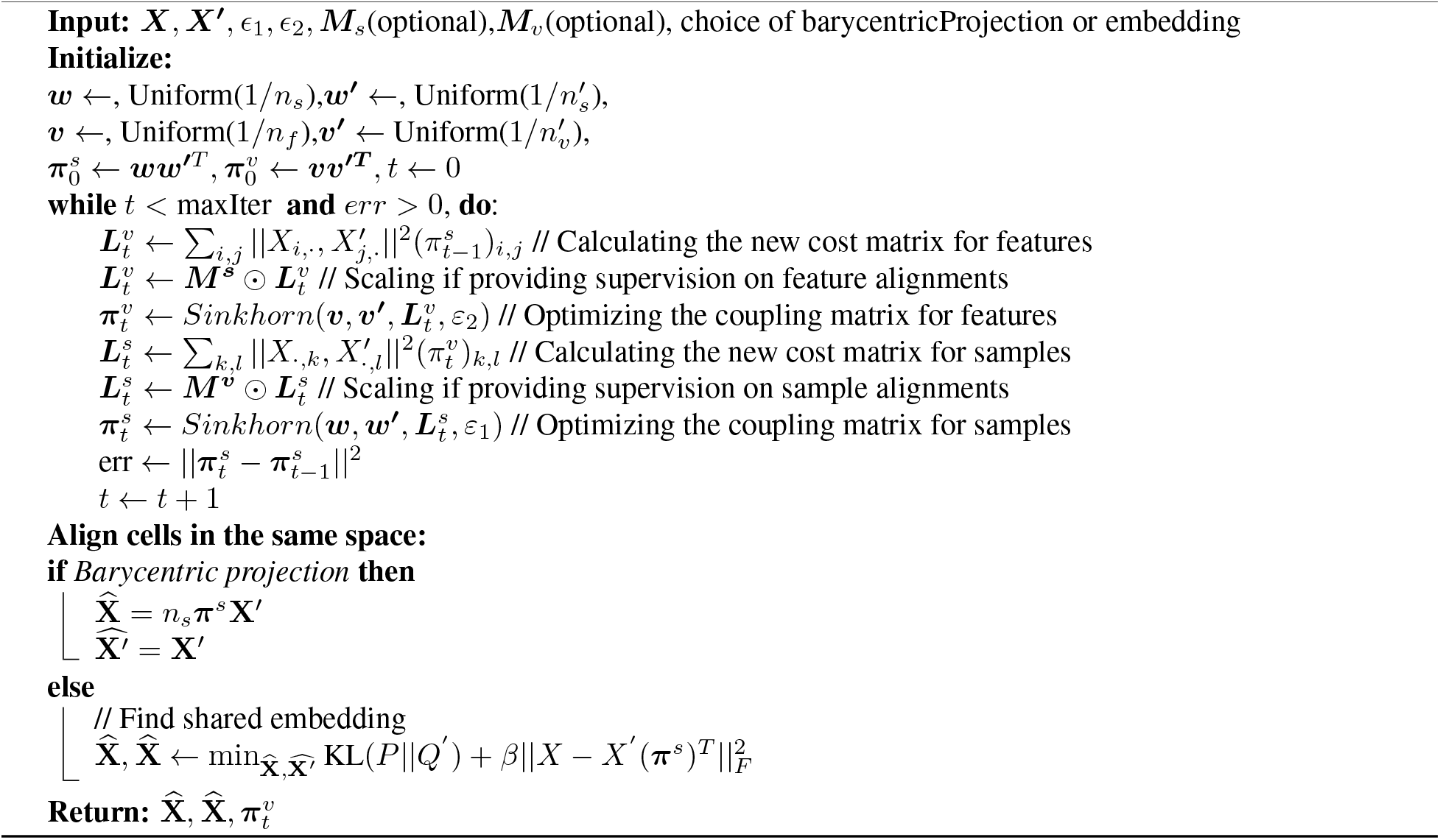
Pseudocode for SCOOTR.

**Algorithm 2:**
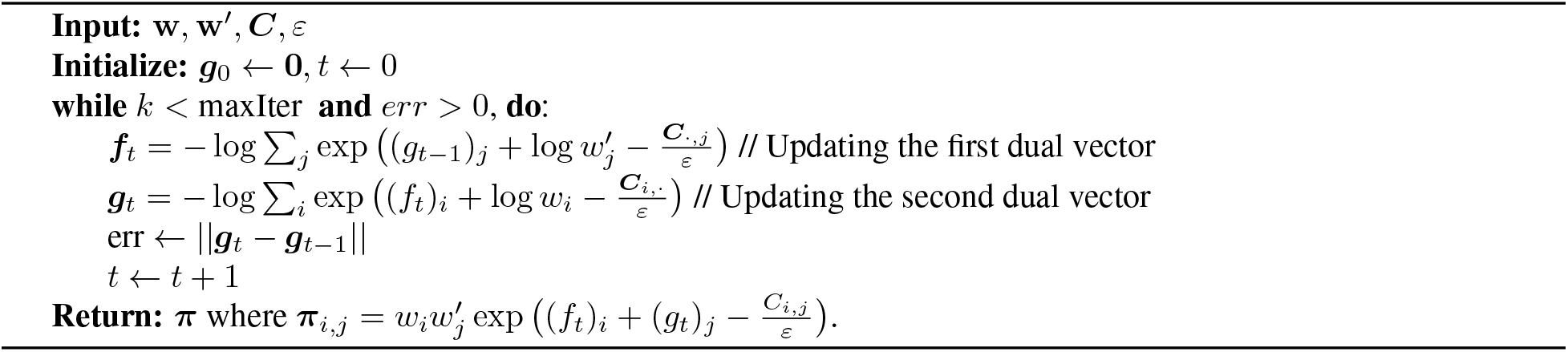
Pseudocode for Sinkhorn iterations.

Here, ⊙ denotes the element-wise multiplication between two matrices, *n_s_* and 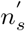 refer to the number of samples in the two input datasets, respectively, and *n_f_* and 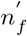 refer to the number of features.

The embedding formulation in the scootr algorithm (which is an alternative to barycentric projection) is based on t-SNE and its details can be found in [30]. This has also been previously used in the other single-cell alignment methods, UnionCom [8], Pamona [10], and SCOTv2 [11]. Briefly, *P*_*j*|*i*_ is the conditional probability that a data point *x_i_* would pick 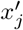 as its neighbor when chosen according a Gaussian distribution centered at *x_i_*. Similarly, *P*_*j*|*i*_ is the same conditional probability, but in the embedding space, and it is computed through a Student-t distribution:

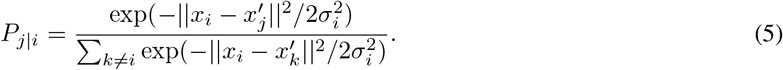

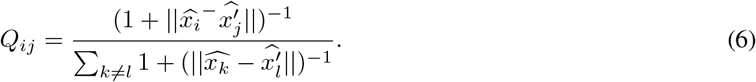

The bandwidth *σ_i_* is chosen according to the density of the data points through a binary search for the value of *σ_i_* that achieves the fixed perplexity value (user can determine), which is computed by averaging *P*_*i*|*j*_ and *P*_*j*|*i*_ to give more weight to outlier points:

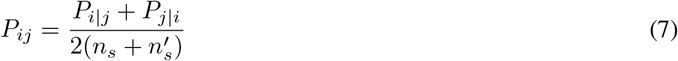

### Hyperparameter Combinations for Baselines

When methods share similar hyperparameters in their formulation (e.g. hyperparameters are dimensionality of the latent space, *p*, for the algorithms that commonly embed datasets; entropic regularization constant, *ϵ*, for methods that employ optimal transport; number of neighbors, *k*, for methods that model single-cell datasets with nearest neighbor graphs). Otherwise, we refer to the publication and the code repository for each method to choose a hyperparameter range.

For SCOT, we tune four hyperparameters: *k* ∈ {20, 30,…, 150}, the number of neighbors in the cell neighborhood graphs, *ϵ* ∈ {5*e* − 4, 3*e* − 4, 1*e* − 4, 7*e* − 3, 5*e* − 3,…, 1*e* − 2}, the entropic regularization coefficient for the optimal transport formulation, *ρ* ∈ {1*e* − 3, 5*e* − 3, 1*e* − 2, 5*e* − 2, 0.1, 1, 10, 100}, *p* ∈ {3, 4, 5, 10, 30, 32}, the output dimension for embedding (and compared it with barycentric projection).

For Pamona, we tune four hyperparameters: *k* ∈ {20, 30,…, 150}, the number of neighbors in the cell neighborhood graphs, *ϵ* ∈ {5*e* − 4, 3*e* − 4, 1*e* − 4, 7*e* − 3, 5*e* − 3,…, 1*e* − 2}, the entropic regularization coefficient for the optimal transport formulation, *λ* ∈ {0.1, 0.5, 1, 5, 10}, the coefficient for the trade-off between aligning corresponding cells and preserving local geometries, and lastly, *p* ∈ {3, 4, 5, 10, 30, 32}, the output dimension for embedding. We choose the ranges for *ϵ* and *k* to be consistent with the corresponding hyperparameters in SCOT and the ranges for the embedding dimensions to be consistent with the recommended values in MMD-MA and UnionCom embeddings.

For UnionCom, we tune the trade-off parameter *β* ∈ {0.1, 1, 5, 10, 15, 20} and the regularization coefficient *ρ* ∈ {0, 0.1, 1, 5, 10, 15, 20} based on the ranges reported by Cao *et al.* in the publication [8]. We additionally tune the maximum neighborhood size permitted in the neighborhood graphs, *k_max_* ∈ {40, 100, 150}, as well as the embedding dimensionality *p* ∈ {3, 4, 5, 10, 30, 32}. The sweep range for hyperparameter *k_max_* is smaller than the other hyperparameters because UnionCom automatically starts from *k* = 2 and goes up to *k_max_* to find the lowest *k* that returns a connected graph to use in the algorithm. Therefore, more refined search is not needed.

For MMD-MA, we choose the weights *λ*_1_ and *λ*_2_ ∈ {1*e* − 2, 5*e* − 3, 1*e* − 3, 5*e* − 4,…, 1*e* − 9}. This range includes the hyperparameter range suggested by Singh *et al* (*λ*_1_, *λ*_2_ ∈ {1*e* − 3, 1*e* − 4, 1*e* − 5, 1*e* − 6, 1*e* − 7}) but extends it further to increase the granularity for the sake of more fair comparison against methods that require a higher number of hyperparameters to test, such as Pamona and UnionCom. Similarly to other methods, we also select the embedding dimensionality from *p* ∈ {3, 4, 5, 10, 30, 32}.

For bindSC, we choose the couple coefficient that assigns weight to the initial gene activity matrix *α* ∈ {0, 0.1, 0.2, 0.9} and the couple coefficient that assigns weight factor to multi-objective function *λ* ∈ {0.1, 0.2, …, 0.9}. Additionally, we choose the number of canonical vectors for the embedding space *K* ∈ {3, 4, 5, 10, 30, 32}.

### Visualizing cell-level alignment of the scGEM dataset

**Figure S1:**
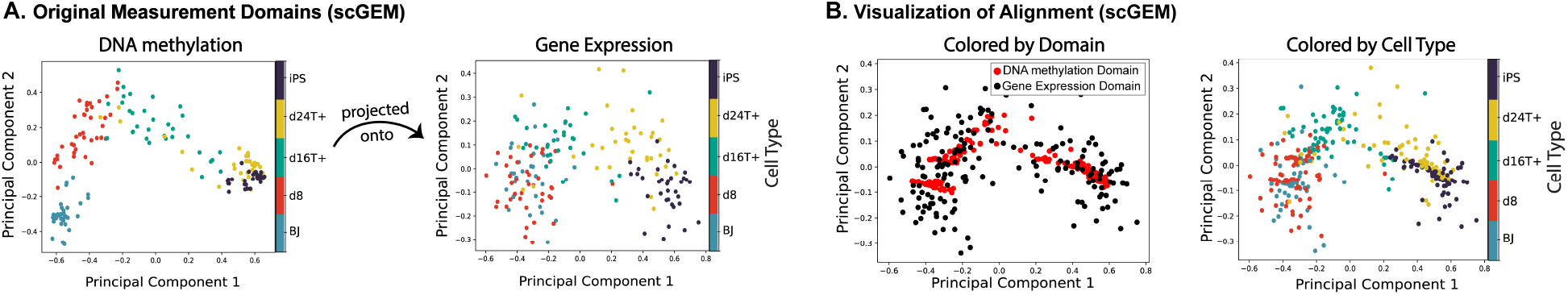
Visualization of scGEM cell-cell alignments. **A**. shows the original measurement domains before alignment, colored by cell type. **B**. plots the aligned datasets upon barycentric projection of the DNA methylation domain onto the gene expression domain with respect to the cell-cell alignment probabilities scootr recovers. On the left-hand side, we color the aligned data points by domain identity, and on the right-hand side, we color them by cell-type identity.

### Approximately self-tuning hyperparameters

**Figure S2:**
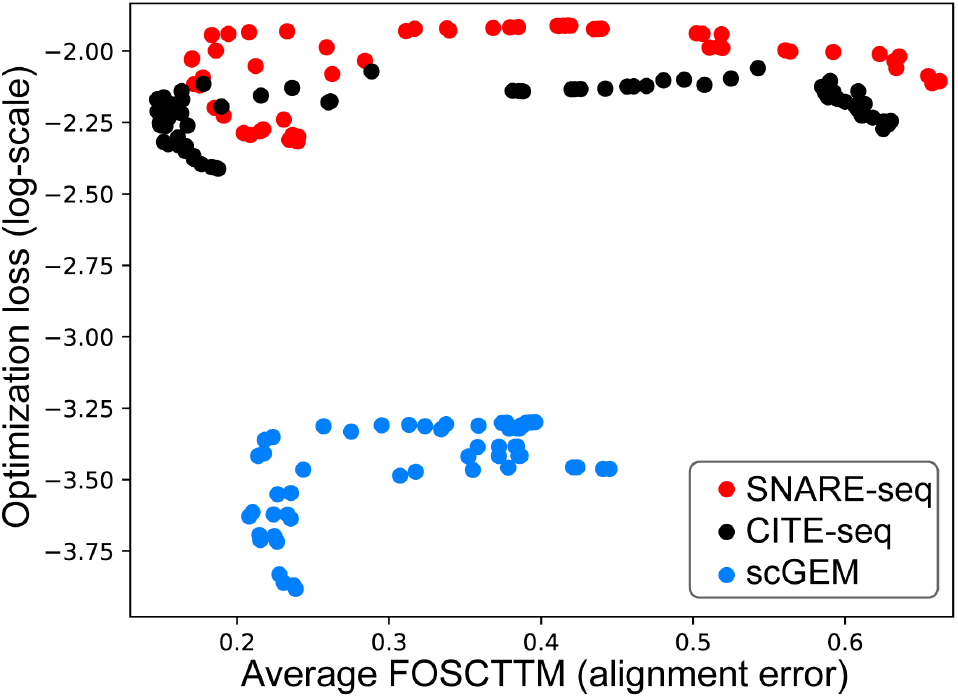
Co-Optimal transport optimization loss (log scale) vs alignment error as measured by average FOS-CTTM. Hyperparameter combinations with lower loss values in optimization tend to yield lower alignment error although it is not a perfect relationship.

### SNARE-seq feature alignments

Figure 3(C) visualizes the top chromatin accessibility feature correspondences for the cell-type marker genes from the gene expression domain. Majority of these correspondences are biologically validated or computationally predicted regulatory relationships, which we discuss here. Firstly, of the ten alignments visualized in this plot, three of them are between marker genes and their chromatin regions. These are (1) *PRAME* and Chr22: 22.520-22.521 Mb region, which is a region upstream of the *PRAME* gene body that is rich with predicted transcriptional factor (TF) binding sites according to the “RepMap Atlas of Regulatory Regions” [**?**] annotations on UCSC Genome Browser (Human hg38 annotations) [26]. Among the predicted TF bindings, many of them are K562-specific predictions, and some of these are known regulators of *PRAME*, such as but not limited to *E2F6*, *HDAC2*, *CTCF* (based on GRNdb database [31] of TF-gene relationships). Additionally, *COL1A2* and *HLA-DRB1* also have recovered correspondences with their own chromosomal region, “Chr7:94.396-94.421 Mb” and “Chr6:32,578-32,579 Mb”, respectively. We observe that *COL1A2* and *PRAME* are also additionally aligned with “Chr1: 58,780 - 58,784 Mb” regions, which correspond to the gene body of *JUN* transcriptional factor. Indeed, *JUN* has been identified as one of the transcriptional factors differentially expressed in the K562 and BJ cells, but more strongly in the latter, according to the original publication that released this dataset [1]. GRNdb also identified *JUN* to be one of the regulators of the *COL1A2* gene. In addition to the chromosomal region of *JUN*, *PRAME* has another region abundant in predicted TF binding sites among its top correspondences: “Chr6: 7.974-7.978 Mb”. This region is annotated with an H3K27Ac mark on the UCSC Genome Browser, and has multiple predicted binding sites of TFs GRNdb identifies as regulators of *PRAME*, such as *IRF1, HDAC2, HOXC6* and POU2AF1. The *HLA-DRB1* gene is also aligned with a chromosomal region rich in GM12878-specific predictions of TF bindings, such as *IRF4*, *IRF8*, *ETV6*, and *CREM*, which GRNdb lists as potential regulators of *HLA-DRB1*. Lastly, even though we couldn’t find a biological relationship reported between the *CLYBL* gene and *EPCAM* gene (marker gene for the H1 cell-line), the chromosomal region in *CLYBL* body where scootr finds a correspondence with *EPCAM* indeed appears to be differentially accessible in H1 cells (and in to a lesser degree, K562) in our dataset.

### Footnotes

1 https://github.com/jw156605/MATCHER

2 https://satijalab.org/seurat/v4.0/weighted_nearest_neighbor_analysis.html

3 https://github.com/PeterZZQ/Symsim2

## Notes

### Competing Interest Statement

The authors have declared no competing interest.

### Summary of Updates

Fixed a few writing errors (tense agreement issues etc) in the second paragraph of the Discussion section.

## References

[1] Song Chen, Blue B. Lake, and Kun Zhang. High-throughput sequencing of the transcriptome and chromatin accessibility in the same cell. Nature Biotechnology, 37(12):1452–1457, 2019.

[2] Marlon Stoeckius, Christoph Hafemeister, William Stephenson, Brian Houck-Loomis, Pratip K Chattopadhyay, Harold Swerdlow, Rahul Satija, and Peter Smibert. Simultaneous epitope and transcriptome measurement in single cells. Nature Methods, 14(9):865–868, 2017.

[3] Stephen J. Clark, Ricard Argelaguet, Chantriolnt-Andreas Kapourani, Thomas M. Stubbs, Heather J. Lee, Celia Alda-Catalinas, Felix Krueger, Guido Sanguinetti, Gavin Kelsey, John C. Marioni, Oliver Stegle, and Wolf Reik. scnmt-seq enables joint profiling of chromatin accessibility dna methylation and transcription in single cells. Nature Communications, 9(1):781, 2018.

[4] Anjun Ma, Adam McDermaid, Jennifer Xu, Yuzhou Chang, and Qin Ma. Integrative methods and practical challenges for single-cell multi-omics. Trends in Biotechnology, 38(9):1007–1022, 2020.

[5] Michael Eisenstein. The secret life of cells. Nature Methods, 17(1):7–10, 2020.

[6] Jie Liu, Yuanhao Huang, Ritambhara Singh, Jean-Philippe Vert, and William Stafford Noble. Jointly Embedding Multiple Single-Cell Omics Measurements. In 19th International Workshop on Algorithms in Bioinformatics (WABI 2019), volume 143, pages 10:1–10:13, 2019.

[7] Ritambhara Singh, Pinar Demetci, Giancarlo Bonora, Vijay Ramani, Choli Lee, He Fang, Zhijun Duan, Xinxian Deng, Jay Shendure, Christine Disteche, and William Stafford Noble. Unsupervised manifold alignment for single-cell multi-omics data. BCB ’20, 2020.

[8] Kai Cao, Xiangqi Bai, Yiguang Hong, and Lin Wan. Unsupervised topological alignment for single-cell multi-omics integration. Bioinformatics, 36(Supplement 1):i48–i56, 2020.

[9] Pinar Demetci, Rebecca Santorella, Björn Sandstede, William Stafford Noble, and Ritambhara Singh. Gromov-wasserstein optimal transport to align single-cell multi-omics data. bioRxiv, 2020.

[10] Kai Cao, Yiguang Hong, and Lin Wan. Manifold alignment for heterogeneous single-cell multi-omics data integration using Pamona. Bioinformatics, 08 2021. btab594.

[11] Pinar Demetci, Rebecca Santorella, Manav Chakravarthy, Bjorn Sandstede, and Ritambhara Singh. Scotv2: Single-cell multiomic alignment with disproportionate cell-type representation. Journal of Computational Biology, 29(11):1213–1228, 2022. PMID: 36251763.

[12] Jinzhuang Dou, Shaoheng Liang, Vakul Mohanty, Qi Miao, Yuefan Huang, Qingnan Liang, Xuesen Cheng, Sangbae Kim, Jongsu Choi, Yumei Li, Li Li, May Daher, Rafet Basar, Katayoun Rezvani, Rui Chen, and Ken Chen. Bi-order multimodal integration of single-cell data. Genome Biology, 23(1):112, 2022.

[13] Gabriel Peyré and Marco Cuturi. Computational optimal transport. Foundations and Trends® in Machine Learning, 11:355–607, 2019.

[14] Facundo Memoli. Gromov wasserstein distances and the metric approach to object matching. Foundations of Computational Mathematics, pages 1–71, 2011.

[15] Gabriel Peyré, Marco Cuturi, and Justin Solomon. Gromov-wasserstein averaging of kernel and distance matrices. In ICML, pages 2664–2672, 2016.

[16] Ievgen Redko, Titouan Vayer, Rémi Flamary, and Nicolas Courty. CO-Optimal Transport. arXiv:2002.03731, 2020.

[17] Marco Cuturi. Sinkhorn distances: Lightspeed computation of optimal transport. In NIPS, pages 2292–2300, 2013.

[18] Luke Zappia, Belinda Phipson, and Alicia Oshlack. Splatter: simulation of single-cell rna sequencing data. Genome biology, 18(1):1–15, 2017.

[19] Lih Feng Cheow, Elise T Courtois, Yuliana Tan, Ramya Viswanathan, Qiaorui Xing, Rui Zhen Tan, Daniel S Q Tan, Paul Robson, Loh Yuin-Han, Stephen R Quake, and William F Burkholder. Single-cell multimodal profiling reveals cellular epigenetic heterogeneity. Nature Methods, 13(10):833–836, 2016.

[20] Kenji Kamimoto, Christy M. Hoffmann, and Samantha A. Morris. Celloracle: Dissecting cell identity via network inference and in silico gene perturbation. bioRxiv, 2020.

[21] Aritra Bhattacherjee, Mohamed Nadhir Djekidel, Renchao Chen, Wenqiang Chen, Luis M. Tuesta, and Yi Zhang. Cell type-specific transcriptional programs in mouse prefrontal cortex during adolescence and addiction. Nature Communications, 10(1):4169, Sep 2019.

[22] Maria Antonietta Tosches, Tracy M. Yamawaki, Robert K. Naumann, Ariel A. Jacobi, Georgi Tushev, and Gilles Laurent. Evolution of pallium, hippocampus, and cortical cell types revealed by single-cell transcriptomics in reptiles. Science, 360(6391):881–888, 2018.

[23] Ziqi Zhang, Chengkai Yang, and Xiuwei Zhang. scDART: integrating unmatched scRNA-seq and scATAC-seq data and learning cross-modality relationship simultaneously. Genome Biology, 23(1):139, 2022.

[24] Xiuwei Zhang, Chenling Xu, and Nir Yosef. Simulating multiple faceted variability in single cell rna sequencing. Nature Communications, 10(1):2611, Jun 2019.

[25] Tim Stuart, Avi Srivastava, Shaista Madad, Caleb A. Lareau, and Rahul Satija. Single-cell chromatin state analysis with signac. Nature Methods, 18(11):1333–1341, Nov 2021.

[26] Jairo Navarro Gonzalez, Ann S Zweig, Matthew L Speir, Daniel Schmelter, Kate R Rosenbloom, Brian J Raney, Conner C Powell, Luis R Nassar, Nathan D Maulding, Christopher M Lee, Brian T Lee, Angie S Hinrichs, Alastair C Fyfe, Jason D Fernandes, Mark Diekhans, Hiram Clawson, Jonathan Casper, Anna Benet-Pagès, Galt P Barber, David Haussler, Robert M Kuhn, Maximilian Haeussler, and W James Kent. The UCSC Genome Browser database: 2021 update. Nucleic Acids Research, 49(D1):D1046–D1057, 11 2020.

[27] Jaime A Castro-Mondragon, Rafael Riudavets-Puig, Ieva Rauluseviciute, Roza Berhanu Lemma, Laura Turchi, Romain Blanc-Mathieu, Jeremy Lucas, Paul Boddie, Aziz Khan, Nicolás Manosalva Pérez, Oriol Fornes, Tiffany Y Leung, Alejandro Aguirre, Fayrouz Hammal, Daniel Schmelter, Damir Baranasic, Benoit Ballester, Albin Sandelin, Boris Lenhard, Klaas Vandepoele, Wyeth W Wasserman, François Parcy, and Anthony Mathelier. JASPAR 2022: the 9th release of the open-access database of transcription factor binding profiles. Nucleic Acids Research, 50(D1):D165–D173, 11 2021.

[28] Fayrouz Hammal, Pierre de Langen, Aurélie Bergon, Fabrice Lopez, and Benoit Ballester. ReMap 2022: a database of Human, Mouse, Drosophila and Arabidopsis regulatory regions from an integrative analysis of DNA-binding sequencing experiments. Nucleic Acids Research, 50(D1):D316–D325, 11 2021.

[29] Cédric Villani. Optimal Transport: Old and New. Grundlehren der mathematischen Wissenschaften. Springer, 2009 edition, September 2008.

[30] Laurens Van der Maaten and Geoffrey Hinton. Visualizing data using t-sne. Journal of machine learning research, 9(11), 2008.

[31] Li Fang, Yunjin Li, Lu Ma, Qiyue Xu, Fei Tan, and Geng Chen. GRNdb: decoding the gene regulatory networks in diverse human and mouse conditions. Nucleic Acids Research, 49(D1):D97–D103, 11 2020.

